# Polygenic transmission disequilibrium confirms that common and rare variation act additively to create risk for autism spectrum disorders

**DOI:** 10.1101/089342

**Authors:** Daniel J. Weiner, Emilie M. Wigdor, Stephan Ripke, Raymond K. Walters, Jack A. Kosmicki, Jakob Grove, Kaitlin E. Samocha, Jacqueline Goldstein, Aysu Okbay, Jonas Bybjerg-Gauholm, Thomas Werge, David M. Hougaard, Jacob Taylor, David Skuse, Bernie Devlin, Richard Anney, Stephan Sanders, Somer Bishop, Preben Bo Mortensen, Anders D. Børglum, George Davey Smith, Mark J. Daly, Elise B. Robinson

**Affiliations:** Analytic and Translational Genetics Unit, Department of Medicine, Massachusetts General Hospital and Harvard Medical School, Boston, Massachusetts, USA.; Stanley Center for Psychiatric Research, Broad Institute of MIT and Harvard, Cambridge, Massachusetts, USA.; Program in Medical and Population Genetics, Broad Institute of MIT and Harvard, Cambridge, Massachusetts, USA.; Department of Psychiatry and Psychotherapy, Charité, Campus Mitte, Berlin, Germany.; Program in Genetics and Genomics, Biological and Biomedical Sciences, Harvard Medical School, Boston, Massachusetts, USA.; Department of Biomedicine (Human Genetics), Aarhus University, Aarhus, Denmark.; Lundbeck Foundation Initiative for Integrative Psychiatric Research, iPSYCH, Copenhagen, Denmark.; Centre for Integrative Sequencing, iSEQ, Aarhus University, Aarhus, Denmark.; Bioinformatics Research Centre, Aarhus University, Aarhus, Denmark.; Department of Complex Trait Genetics, Center for Neurogenomics and Cognitive Research, Vrije Universiteit Amsterdam, Amsterdam, 1081 HV, The Netherlands.; Danish Center for Neonatal Screening, Department for Congenital Disorders, Statens Serum Institut, Copenhagen, Denmark.; Institute of Biological Psychiatry, Mental Health Center Sct. Hans, Mental Health Services Copenhagen, Copenhagen, Denmark.; Department of Clinical Medicine, University of Copenhagen, Copenhagen, Denmark.; Department of Psychiatry, Brigham and Women’s Hospital, Boston, Massachusetts, USA.; Behavioural Sciences Unit, Institute of Child Health, University College London, London, UK.; Department of Psychiatry, University of Pittsburgh School of Medicine, Pittsburgh, Pennsylvania, USA.; Division of Psychological Medicine and Clinical Neurosciences, Cardiff University, Cardiff, Wales, UK.; Department of Psychiatry, University of California, San Francisco, San Francisco, California, USA.; National Centre for Register-based Research, University of Aarhus, Aarhus, Denmark.; Medical Research Council Integrative Epidemiology Unit, University of Bristol, Bristol, UK.; iPSYCH, The Lundbeck Foundation Initiative for Integrative Psychiatric Research, Denmark.; Center for Neonatal Screening, Department for Congenital Disorders, Statens Serum Institut, Copenhagen, Denmark.; Institute of Biological Psychiatry, MHC Sct. Hans, Mental Health Services Copenhagen, Roskilde, Denmark.; iSEQ, Centre for Integrative Sequencing, Aarhus University, Aarhus, Denmark.; Department of Biomedicine - Human Genetics, Aarhus University, Aarhus, Denmark.; Psychosis Research Unit, Aarhus University Hospital, Risskov, Denmark.; Mental Health Services in the Capital Region of Denmark, Mental Health Center Copenhagen, University of Copenhagen, Copenhagen, Denmark.; Analytic and Translational Genetics Unit, Dept. of Medicine, Massachusetts General Hospital and Harvard Medical School, Boston, MA 02114, USA.; Stanley Center for Psychiatric Research and Program in Medical and Population Genetic, Broad Institute of Harvard and MIT, Cambridge, MA 02142, USA.; MRC Centre for Neuropsychiatric Genetics & Genomics, Cardiff University, Cardiff, CF24 4HQ, UK.; Dept. of Psychiatry, University of Pittsburgh School of Medicine, Pittsburgh, PA 15213, USA.; Dept. of Psychiatry, Harvard Medical School, Boston, MA 02115, USA.; Queensland Institute of Medical Research, Brisbane, QLD, 4006, Australia.; Dept. of Psychiatry, University of California San Francisco, San Francisco, CA 94143, USA.; Inst. Human Genetics, University of California San Francisco, San Francisco, CA 94143, USA.; Dept. of Pediatrics, University of Alberta, Edmonton, AB, T6G 1C9, Canada.; Division of Genetics, Children’s Hospital Boston, Harvard Medical School, Boston, MA 02115, USA.; School of Education, University of Birmingham, Birmingham, B15 2TT, UK.; Dept. of Medicine, University of Washington, Seattle, WA 98195, USA.; Dept. of Biostatistics, University of Washington, Seattle, WA 98195, USA.; Dept. of Psychiatry, Carver College of Medicine, Iowa City, IA 52242, USA.; Dept. of Child and Adolescent Psychiatry, Psychosomatics and Psychotherapy, JW Goethe University Frankfurt, Frankfurt am Main, 60528, Germany.; Program in Genetics and Genomics, Harvard Medical School, Boston, MA 02115, USA.; Howard Hughes Medical Institute, Harvard Medical School, Boston, MA 02115, USA.; Dept. of Pediatrics, Harvard Medical School, Boston, MA 02115, USA.; Dept. of Neurology, Harvard Medical School, Boston, MA 02115, USA.; Dept. of Psychiatry, University of Oxford and Warneford Hospital, Oxford, OX3 7JX, UK.; Dept. of Psychiatry, Brain Center Rudolf Magnus, University Medical Center Utrecht, Utrecht, 3584 CG, The Netherlands.; Battelle Center for Mathematical Medicine, The Research Institute at Nationwide Children’s Hospital, Columbus, OH 43205, USA.; Instituto Nacional de Saœde Dr. Ricardo Jorge, Lisboa, 1600, Portugal.; Center for Biodiversity, Functional and Integrative Genomics, Campus da FCUL, Lisboa, 1649, Portugal.; Dept. of Psychiatry and Behavioral Neurosciences, McMaster University, Hamilton, ON, L8S 4L8, Canada.; Dept. of Psychiatry, University of Toronto, ON, M5T 1R8, Canada.; Karolinska Institutet, Solna, SE-171 77, Sweden.; deCODE Genetics, Reykjavik, IS-101, Iceland.; Seaver Autism Center for Research and Treatment, Icahn School of Medicine at Mount Sinai, New York, NY 10029, USA.; Dept. of Psychiatry, Icahn School of Medicine at Mount Sinai, New York, NY 10029, USA.; Dept. of Psychiatry, Rush University Medical Center, Chicago, IL 60612, USA.; Dept. of Psychiatry and Drug Addiction, Tbilisi State Medical University, Tbilisi, 0186, Georgia.; The Centre for Applied Genomics, The Hospital for Sick Children, Toronto, ON, M5G 1L4, Canada.; McLaughlin Centre, University of Toronto, Toronto, ON, M5G 0A4, Canada.; Dept. of Molecular Genetics, University of Toronto, Toronto, ON, M5S 1A8, Canada.; Dept. of Pathology and Laboratory Medicine, University of Pennsylvania, Philadelphia, PA 19102, USA.; State Diagnostic and Counseling Centre, Kopavogur, IS-201, Iceland.; Montreal Neurological Institute, Dept of Neurology and Neurosurgery, McGill University, Montreal, QC, H3A 2B4, Canada.; Centre d’Etudes et de Recherches en Psychopathologie, Toulouse University, Toulouse, 31058, France.; Dept. of Computational Biology, Carnegie Mellon University, Pittsburgh, PA 15213, USA.; Dept. of Statistics, Carnegie Mellon University, Pittsburgh, PA 15213, USA.; Autism Research Unit, The Hospital for Sick Children, Toronto, ON, M5G 1L4, Canada.; Sanger Institute, Hinxton, CB10 1SA, UK.; National Childrens Research Centre, Our Lady’s Hospital Crumlin, Dublin, D12, Ireland.; Academic Centre on Rare Diseases, University College Dublin, Dublin, D4, Ireland.; University of North Carolina, Chapel Hill, NC 27599, USA.; Dept. of Genetics and Genomic Sciences, Icahn School of Medicine at Mount Sinai, New York, NY 10029, USA.; The Mindich Child Health and Development Institute, Icahn School of Medicine at Mount Sinai, New York, NY 10029, USA.; The Icahn Institute for Genomics and Multiscale Biology, Icahn School of Medicine at Mount Sinai, New York, NY 10029, USA.; Friedman Brain Institute, Icahn School of Medicine at Mount Sinai, New York, NY 10029, USA.; The John P. Hussman Institute for Human Genomics, University of Miami, Miami, FL 33101, USA.; Institute of Mental Health and Medical Faculty, University of Belgrade, Belgrade, 11 000, Serbia.; iPSYCH, Lundbeck Foundation Initiative for Integrative Psychiatric Research, Aarhus, Denmark.; National Centre for Register-based Research, Aarhus University, Aarhus, Denmark.; Centre for Integrated Register-based Research, Aarhus University, Aarhus, Denmark.; Dalla Lana School of Public Health, Toronto, ON, M5T 3M7, Canada.; Institute of Neuroscience, Newcastle University, Newcastle Upon Tyne, NE2 4HH, UK.; Institue of Health and Science, Newcastle University, Newcastle Upon Tyne, NE2 4AX, UK.; Wellcome Trust Centre for Human Genetics, Oxford University, Oxford, OX3 7BN, UK.; Unidade de Neurodesenvolvimento e Autismo do Serviço do Centro de Desenvolvimento da Criança and Centro de Investigação e Formação Clinica, Pediatric Hospital, Centro Hospitalar e Universitário de Coimbra, Coimbra, 3041-80, Portugal.; University Clinic of Pediatrics and Institute for Biomedical Imaging and Life Science, Faculty of Medicine, University of Coimbra, Coimbra, 3041-80, Portugal.; Institute of Psychiatric Research, Dept. of Psychiatry, Indiana University School of Medicine, Indianapolis, IN 46202, USA.; Dept. of Medical and Molecular Genetics, Indiana University School of Medicine, Indianapolis, IN 46202, USA.; Program in Medical Neuroscience, Indiana University School of Medicine, Indianapolis, IN 46202, USA.; Programs on Neurogenetics, Yale University School of Medicine, New Haven, CT 06520, USA.; Dept. of Psychiatry and Human Behaviour, Brown University, Providence, RI 02912, USA.; Tufts University, Boston, MA 02155, USA.; Dept. of Psychiatry, Trinity College Dublin, Dublin, D8, Ireland.; Dept. of Psychiatry, University of Utah, Salt Lake City, UT 84108, USA.; Dept. of Pediatrics, Vanderbilt University, Nashville, TN 37232, USA.; iSEQ, Centre for Integrative Sequencing, Aarhus University, Aarhus, DK-8000, Denmark.; Dept. of Biomedicine - Human Genetics, Aarhus University, Aarhus, DK-8000, Denmark.; Dept. of Child, Adolescent Psychiatry and Medical-Social Rehabilitation, Ukrainian Research Institute of Social Forensic Psychiatry and Drug Abuse, Kyiv, 04080, Ukraine.; Dept. of Pediatrics and Human Genetics, University of Michigan, Ann Arbor, MI 48109, USA.; Yale Center for Genomic Analysis, Yale University School of Medicine, New Haven, CT 06516, USA.; Dept. of Child and Adolescent Psychiatry, National University Hospital, Reykjavik, IS-101, Iceland.; Dept. of Pharmacy and Biotechnology, University of Bologna, Bologna, 40126, Italy.; Center for Autism Research and Treatment, Semel Institute, David Geffen School of Medicine at University of California Los Angeles, Los Angeles, CA 90095, USA.; Program in Neurogenetics, Dept. of Neurology, David Geffen School of Medicine, University of California, Los Angeles, Los Angeles, CA 90095, USA.; Center for Neurobehavioral Genetics, Semel Institute, David Geffen School of Medicine, University of California, Los Angeles, Los Angeles, CA 90095, USA.; Dept. of Psychiatry, Weill Cornell Medical College, Cornell University, New York, NY 10065, USA.; Dept. of Pediatrics, Keck School of Medicine, University of Southern California, Los Angeles, CA 90027, USA.; Autism & Developmental Medicine Institute, Geisinger Health System, Danville, PA 17837, USA.; Chief Scientific Officer, Geisinger Health System, Danville, PA 17837, USA.; FondaMental Foundation, Créteil, 94000, France.; INSERM U955, Paris, 94010, France.; Faculté de Médecine, Université Paris Est, Créteil, 94000, France.; Dept. of Psychiatry, Henri Mondor-Albert Chenevier Hospital, Assistance Publique – HÔpitaux de Paris, Créteil, 94000, France.; Dept. of Epidemiology, Johns Hopkins Bloomberg School of Public Health, Baltimore, MD 21205, USA.; Division of Molecular Genome Analysis and Working Group Cancer Genome Research, Deutsches Krebsforschungszentrum, Heidelberg, D-69120, Germany.; Institute for Juvenile Research, Dept. of Psychiatry, University of Illinois at Chicago, Chicago, IL 60612, USA.; Institute of Translational Neuroscience and Dept. of Psychiatry, University of Minnesota, Minneapolis, MN 55454, USA.; Dept. of Public Health Sciences, School of Medicine, University of California Davis, Davis, CA 95616, USA.; The MIND Institute, School of Medicine, University of California Davis, Davis, CA 95817, USA.; Center for Neonatal Screening, Dept. for Congenital Disorders, Statens Serum Institut, Copenhagen, DK-2300, Denmark.; Dept. of Molecular Physiology & Biophysics, Vanderbilt University, Nashville, TN 37232, USA.; Manchester Academic Health Sciences Centre, Manchester, M13 9NT, UK.; Institute of Brain, Behaviour, and Mental Health, University of Manchester, Manchester, M13 9PT, UK.; Centre for Medical Genetics, Our Lady’s Hospital Crumlin, Dublin, D12, Ireland.; Gillberg Neuropsychiatry Centre, University of Gothenburg, Gothenburg, S-405 30, Sweden.; Dept. of Psychiatry and Institute for Development and Disability, Oregon Health & Science University, Portland, OR 97239, USA.; Division of Child and Adolescent Psychiatry, Dept. of Psychiatry, Miller School of Medicine, University of Miami, Miami, FL 33136, USA.; Memorial University of Newfoundland, St. John’s, NL, A1B 3X9, Canada.; Dept. of Mental Health, Johns Hopkins Bloomberg School of Public Health, Baltimore, MD 21205, USA.; Human Genetics and Cognitive Functions Unit, Institut Pasteur, Paris, 75015, France.; Centre National de la Recherche Scientifique URA 2182 Institut Pasteur, Paris, 75724, France.; Dept. of Child and Adolescent Psychiatry, Robert Debré Hospital, Assistance Publique – HÔpitaux de Paris, Paris, 75019, France.; Duke Center for Autism and Brain Developments, Duke University School of Medicine, Durham, NC 27705, USA.; Duke Institute for Brain Sciences, Duke University School of Medicine, Durham, NC 27708, USA.; Temple Street Children’s University Hospital, Dublin, D1, Ireland.; Dept. of Molecular and Human Genetics, Baylor College of Medicine, Houston, TX 77030, USA.; Dept. of Psychiatry, David Geffen School of Medicine at University of California Los Angeles, Los Angeles, CA 90095, USA.; Dept. of Human Genetics, David Geffen School of Medicine at University of California Los Angeles, Los Angeles, CA 90095, USA.; University Paris Diderot, Sorbonne Paris Cité, Paris, 75013, France.; Institute of Psychiatry, Kings College London, London, SE5 8AF, UK.; South London & Maudsley Biomedical Research Centre for Mental Health, London, SE5 8AF, UK.; Dept. of Women’s and Children’s Health, Center of Neurodevelopmental Disorders, Karolinska Institutet, Stockholm, SE-113 30, Sweden.; Child and Adolescent Psychiatry, Center for Psychiatry Research, Stockholm County Council, Stockholm, SE-171 77, Sweden.; INSERM U1130, Paris, 75005, France.; CNRS UMR 8246, Paris, 75005, France.; Sorbonne Universités, UPMC Univ Paris 6, Neuroscience Paris Seine, Paris, 75005, France.; Dept. of Psychiatry and Behavioral Sciences, University of Washington, Seattle, WA 98195, USA.; Stella Maris Institute for Child and Adolescent Neuropsychiatr, Pisa, 56018, Italy.; Paediatric Neurodisability, King’s Health Partners, Kings College London, London, SE1 7EH, UK.; Mental Health and Addictions Research Unit, University of British Colombia, Vancouver, BC, V5Z 4H4, Canada.; McKusick-Nathans Institute of Genetic Medicine, Johns Hopkins University, Baltimore, MD 21218, USA.; Bloorview Research Institute, University of Toronto, Toronto, ON, M4G 1R8, Canada.; Dept. of Psychiatry, School of Medicine, University of California Davis, Davis, CA 95817, USA.; Dept. of Behavioural Sciences, School of Medicine, University of California Davis, Davis, CA 95817, USA.; Dept. of Neuroscience, Icahn School of Medicine at Mount Sinai, New York, NY 10029, USA.; The Center for Applied Genomics and Division of Human Genetics, ChildrenÕs Hospital of Philadelphia, University of Pennsylvania School of Medicine, Philadelphia, PA 19104, USA.; Dept of Pediatrics, University of Pennsylvania, Philadelphia, PA 19104, USA.; Dept. of Psychiatry, Stanford University, Stanford, CA 94305, USA.; Maine Medical Center Research Institute, Portland, ME 04074, USA.; Center for Human Genetics Research, Vanderbilt University, Nashville, TN 37232, USA.

## Abstract

Autism spectrum disorder (ASD) risk is influenced by both common polygenic and *de novo* variation. The purpose of this analysis was to clarify the influence of common polygenic risk for ASDs and to identify subgroups of cases, including those with strong acting *de novo* variants, in which different types of polygenic risk are relevant. To do so, we extend the transmission disequilibrium approach to encompass polygenic risk scores, and introduce the polygenic transmission disequilibrium test. Using data from more than 6,400 children with ASDs and 15,000 of their family members, we show that polygenic risk for ASDs, schizophrenia, and greater educational attainment is over transmitted to children with ASDs in two independent samples, but not to their unaffected siblings. These findings hold independent of proband IQ. We find that common polygenic variation contributes additively to risk in ASD cases that carry a very strong acting *de novo* variant. Lastly, we find evidence that elements of polygenic risk are independent and differ in their relationship with proband phenotype. These results confirm that ASDs’ genetic influences are highly additive and suggest that they create risk through at least partially distinct etiologic pathways.

Risk for autism spectrum disorders (ASDs) is strongly genetically influenced, and reflects several types of genetic variation^1-7^. Common polygenic variation, distributed across the genome, accounts for at least 20% of ASD liability^2,5,8,9^. *De novo* single nucleotide and copy number variants can strongly affect the individuals who carry them^1,3,4^, but account for less liability at a population level (< 10%)^2^. Over the last several years, the common polygenic and *de novo* influences on ASD risk have been increasingly well characterized, particularly in terms of the distribution of their phenotypic effects. Most consistently, ASD-associated *de novo* variants have been strongly linked to intellectual disability, as well as other indicators of global neurodevelopmental impact (e.g., seizures; motor delay)^1,10^. Indeed ASD-associated *de novo* mutations that yield protein truncations are far more commonly observed in global developmental delay than in autism itself^11^.

Recent studies have suggested that the common polygenic component of ASDs has a different, perhaps surprising, relationship with cognition. Polygenic risk for ASDs has been positively associated with intelligence and educational attainment in several reports^6,12-14^. In other words, in the general population, greater common variant risk for ASDs is associated with higher IQ. These findings are difficult to interpret – on average, IQ in ASD is at least a standard deviation below the population mean^1,15^. Further, ASDs share approximately 25% of their common variant influences with schizophrenia, and schizophrenia itself shows a negative genetic correlation with IQ^6,14^. While genetic correlation analyses are not expected to be transitive, this particularly complex network of common variant associations has led to concerns about confounding resulting from ascertainment and case heterogeneity.

Here we attempt to clarify the influence of common variant risk for ASDs, and to better understand the subgroups of ASD individuals for whom polygenic variation is risk contributing. We thus sought to employ a robust family-based design that would be immune to many of the potential confounders that can arise in attempting to construct a well-matched case-control comparison. To advance this analysis we extended the transmission disequilibrium approach to encompass polygenic risk scores; we call the resulting methodology the polygenic transmission disequilibrium test (pTDT). Using pTDT we then show that common polygenic predictors of (i) ASDs, (ii) schizophrenia, and (iii) years of educational attainment are unambiguously associated with ASD risk, independent of the presence of intellectual disability in cases. We find that common polygenic variation still contributes to ASD risk in cases that carry a very strong acting *de novo* event. Lastly, we find that the three aforementioned polygenic risk factors have independent and distinct effects on phenotypic heterogeneity in ASD, suggesting that components of common polygenic variation also behave additively and operate through at least partially distinct etiologic pathways.

## Polygenic Transmission Disequilibrium

Children are expected to inherit half of their parents’ risk alleles for a trait. This expectation forms the basis of several commonly used tests of genetic association. The classic transmission disequilibrium test, for example, examines the frequency with which single genetic variants are transmitted from parents to their children^16^. Variants transmitted significantly more than half of the time from unaffected parents to children affected with some trait or condition are nominated for association with that trait – only the ascertainment on a trait in the offspring and the association of the allele with the trait introduces deviation from the 50-50 chance of inheriting either allele from a heterozygous parent. Transmission disequilibrium tests have several convenient properties. First, they are immune to confounding by ancestry. They are also less vulnerable to bias from other potential differences between cases and controls, such as socioeconomic background or other factors commonly related to case ascertainment since the ‘controls’ are, in effect, the perfectly matched untransmitted chromosomes.

In this study, we extend the transmission disequilibrium test to polygenic risk scores and introduce the polygenic transmission disequilibrium approach. A polygenic risk score (PRS) provides a quantitative measure of an individual’s genome-wide common variant predisposition (or “risk”) for a trait (**Online Methods: *Polygenic Risk Scoring***). PRS are normally distributed in the population, which means that some degree of common variant risk for complex traits like ASDs is present in all of us. Since a child has a 50% chance of inheriting either allele from a heterozygous parent, it is algebraically defined that the expected value of a child’s PRS for any trait will be equal to the average of their parents’ PRS (mid-parent PRS). To use height as a conceptual illustration, one expects the average PRS for a group of a children to be the sum of their mothers’ PRS for height and their fathers’ PRS for height, divided by two (Figure 1a). This corresponds to the phenotypic observation that children’s height averages to the sex-standardized mean height of their parents^17^. Of course, children are often taller or shorter than expected; if one were to select a subgroup of children each several inches taller than expected, their average height PRS should exceed that of their parents (Figure 1b). A statistically significant deviation between (i) the mid-parent PRS distribution and (ii) the mean of the child PRS distribution would provide strong evidence that common polygenic influences contributed to high height in the offspring. We refer to this deviation as polygenic transmission disequilibrium.

**Figure 1.**
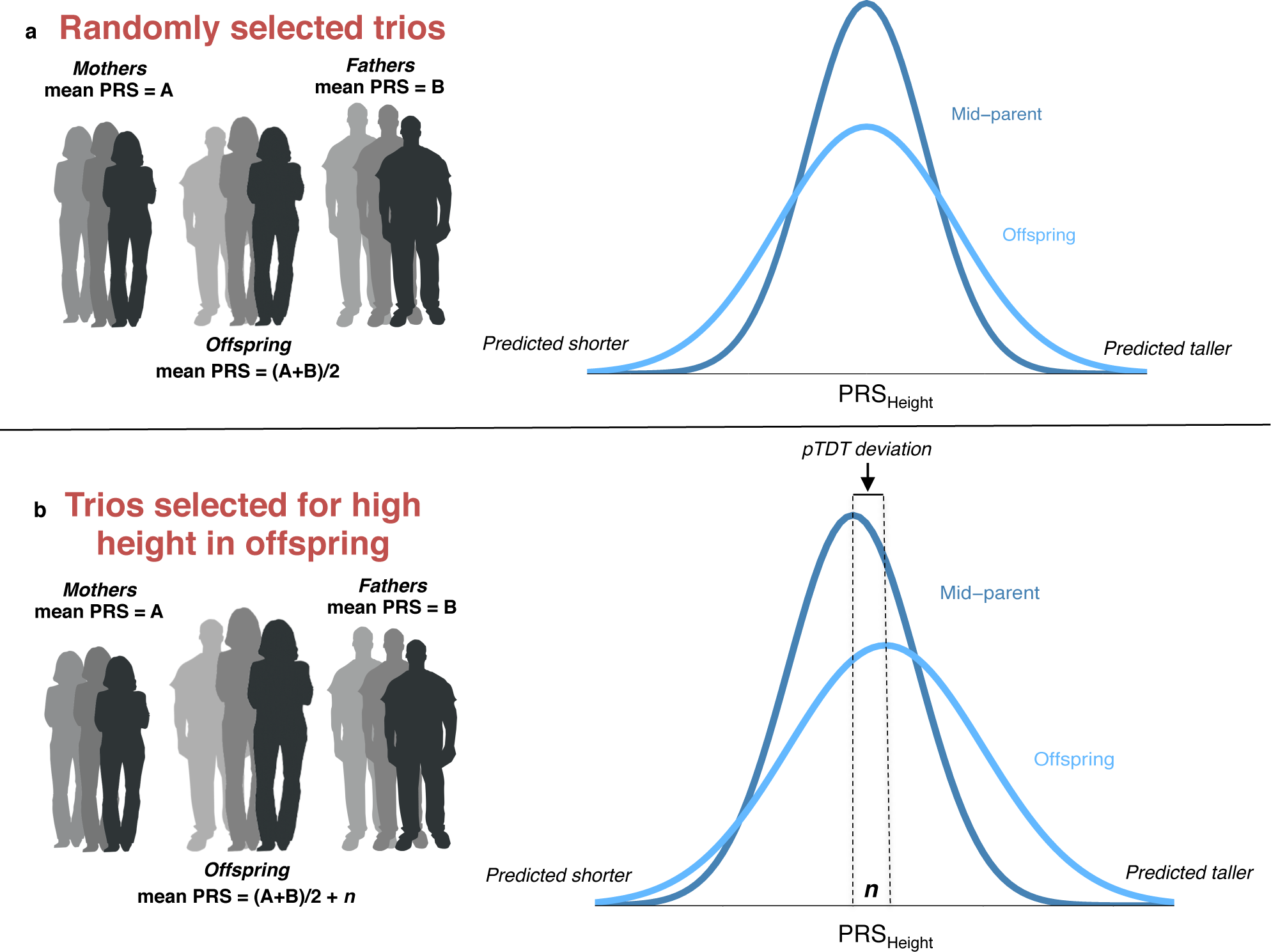
(**a**) In a randomly selected cohort of parent-child trios, a child’s genetic predisposition for height (polygenic risk score, or PRS) is expected to be the average of their parents’ PRS (mid-parent PRS). This corresponds to the phenotypic observation that children’s height is expected to be the sex-standardized average height of their parents. (**b**) In a cohort of trios selected for very high height in the offspring, we correspondingly expect offspring PRS to exceed mid-parent PRS. The difference between the mean of the offspring PRS distribution and mod-parent PRS distribution, *n*, we refer to as polygenic transmission disequilibrium.

Similar logic can be applied to neuropsychiatric disorders such as ASDs^18^. Individuals diagnosed with ASDs should, on average, have more common variant risk for ASDs than their unaffected family members^9^. By comparing ASD individuals’ PRS for various traits against those of their unaffected parents, one can unambiguously associate specific types of common variant risk (e.g., polygenic risk for schizophrenia, educational attainment) with ASDs, and query the subsets of cases in which those risk factors are most relevant.

We used two independent ASD family cohorts to examine transmission of polygenic risk (**Supplementary Table 1; Online Methods: *Sample Description***). The Simons Simplex Collection (SSC) is a resource of more than 2,500 families with a single child diagnosed with ASD^19^. No other family members to the level of first cousins had an ASD diagnosis. Genotype data were available for the parents, the affected child, and an unaffected sibling for 2,091 SSC families (quad families). An additional 493 SSC families had available data for the parents and the affected child alone (trio families). Most genotyped individuals in SSC were exome sequenced in previous studies (89.0%). Our independent Psychiatric Genomics Consortium ASD (PGC ASD) sample consisted of 3,870 genotyped parent-child trios from the Psychiatric Genomics Consortium Autism Group (**Supplementary Table 2**). The PGC ASD cohort described here does not include individuals from the SSC, and the data set does not include exome sequence information.

Using a standard approach^20^, we calculated common polygenic risk for ASDs, educational attainment (EA) and schizophrenia (SCZ) for all genotyped family members in the SSC and PGC ASD datasets (**Online Methods: *Polygenic Risk Scoring***). To do so, we used summary statistics from the largest genome-wide association studies of each phenotype (**Supplementary Table 3).** All discovery GWAS (including ASD) were independent, in that they did not include any SSC or PGC ASD individuals. Using the calculated polygenic risk scores, we first examined properties of their distribution in the parents. Analyzing the SSC and PGC ASD cohorts separately, we did not observe any consistent correlations between the parents’ PRS, for any pair of PRS, and based on that no PRS-based evidence of assortative mating (**Supplementary Method 1**, **Supplementary Tables 4a-b**). We also found no evidence that the mothers and fathers of children with ASD differ with regard to the common variant risk that they carry (**Supplementary Methods 2a, Supplementary Table 5**). Lastly, there was no evidence that mid-parent polygenic risk differs by proband sex (**Supplementary Methods 2b, Supplementary Table 6**).

The pTDT is a t-test asking whether the mean of the offspring PRS distribution is consistent with its parentally-derived expected value (**Online Methods: *pTDT***). In brief, for each trio, one averages the parent PRS for a given trait, ASD for example, and then subtracts that value from the proband’s PRS for the same trait. To standardize and improve interpretability, we divide the resulting difference by the standard deviation of the (observed) mid-parent distribution. This yields the estimated pTDT deviation, and the pTDT then tests whether the average pTDT deviation across all offspring differs from zero. A mean pTDT deviation of 0.1 for the ASD PRS, for example, would indicate that the offspring’s ASD PRS is on average 0.1 standard deviations higher than that of their parents.

For each of ASD, SCZ, and EA, we calculated the average pTDT deviation for each affected child in SSC and PGC ASD, as well as for the unaffected siblings in the SSC. The primary pTDT results are shown in Figure 2a. In both the SSC and PGC ASD samples, polygenic risk for each of ASD, EA, and SCZ was significantly over transmitted to affected children (P < 1E-06 for all comparisons), but not to SSC unaffected siblings (P > 0.05 for all comparisons; Figure 2a). This means that common polygenic risk defined in GWAS studies of ASD, SCZ, and greater EA are each unambiguously associated with autism spectrum disorders. The results did not change when the SSC and PGC ASD samples were restricted to families of European ancestry (**Supplementary Methods 3**; **Supplementary Table 7a**). We repeated the analysis in SSC and PGC ASD using a PRS for body mass index (BMI) – a polygenic risk category unassociated with ASD^6^ – as a negative control (**Online Methods: *Polygenic Risk Scoring***). As expected, we found that neither ASD probands nor unaffected siblings over inherited BMI PRS (P > 0.05 for all comparisons) (**Supplementary Table 8**).

**Figure 2.**
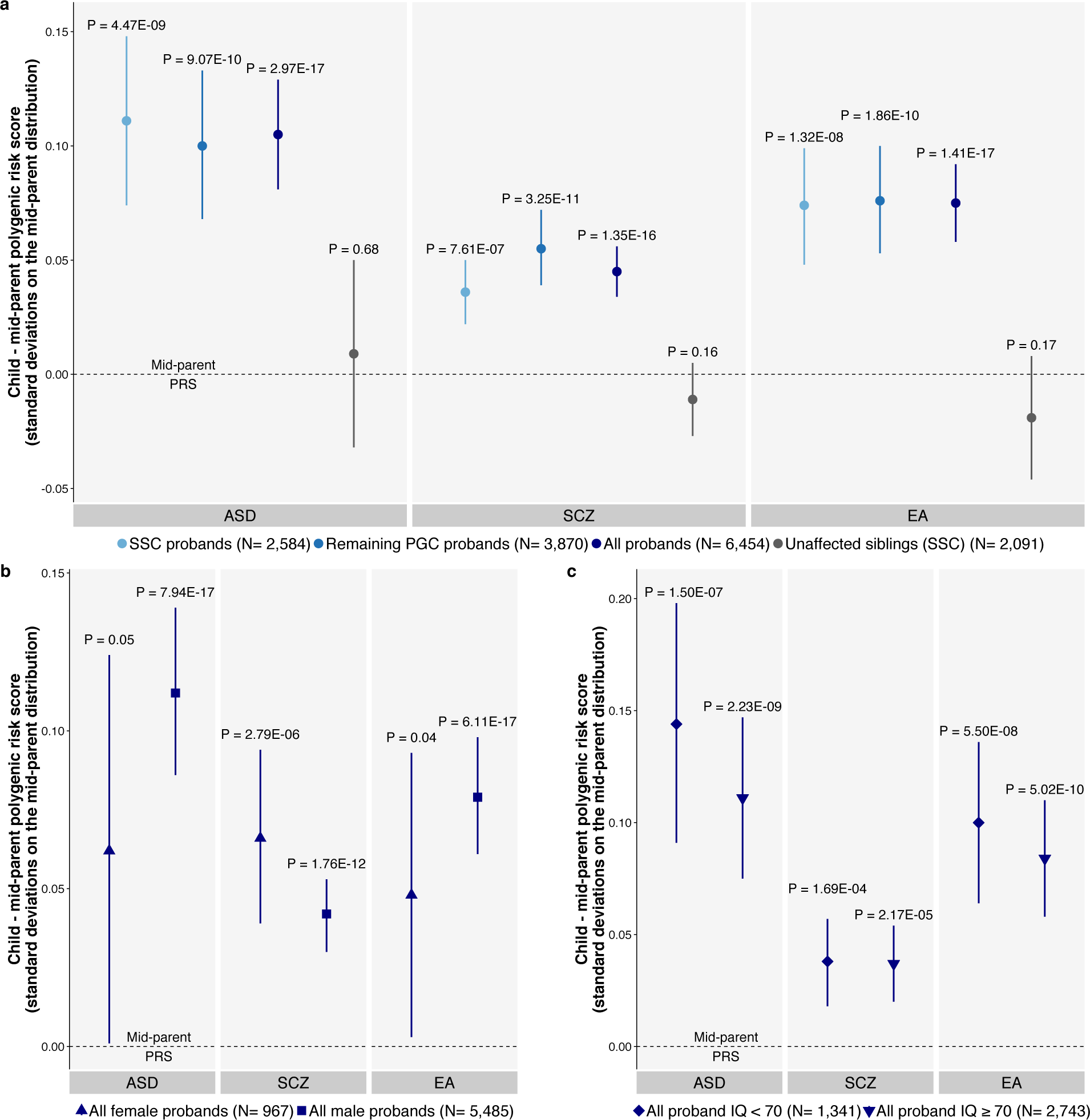
ASD probands over inherit polygenic risk for ASD, schizophrenia, and educational attainment. Transmission is shown in terms of standard deviations on the average parent distribution α 1.96 SE (95% confidence intervals). P-values denote the probability that the mean of the pTDT deviation distribution is 0 (one-sample t-test). (**a**) ASD probands over inherit ASD associated polygenic risk in the Simon Simplex Collection (SSC), Psychiatric Genomics Consortium Autism Group (PGC ASD), and combined cohorts. Unaffected siblings in SSC do not over inherit ASD associated polygenic risk. (**b**) Both male and female probands over inherit ASD associated polygenic risk in the combined cohort. (**c**) ASD probands with and without intellectual disability (full-scale IQ < 70) over inherit ASD associated polygenic risk in the combined cohort.

The degree of polygenic over transmission did not differ between SSC and PGC ASD probands for any comparison (P > 0.05 for all comparisons**; Supplementary Table 7b**), so we combined the samples to improve the power of subsequent subgroup analyses. Using the combined data, each PRS is over transmitted to both male and female probands (P < 0.05 for all comparisons; Figure 2b). In Figure 2c, we also see that each of the three PRS are significantly, and equally, over transmitted to ASD cases with measured IQs in the intellectual disability range (IQ < 70, (**Online Methods: *pTDT***; **Supplementary Tables 9a-b**) when compared with those without intellectual disability. Most probands in the intellectual disability group do not have an observed genetic event that might explain their low IQ, for example a *de novo* mutation from an ASD-associated class, and repeating the analysis in the low IQ SSC probands with *de novo* carriers removed did not change the observed associations (**Supplementary Table 10; Online Methods: De novo *variant analyses***).

## Common and rare variant additivity

In order to best interrogate additivity between common polygenic and rare, strong acting variation, we defined a group of *de novo* mutations of large effect (**Supplementary Table 11**). We previously identified a subclass of *de novo* protein truncating variants (contributing PTVs) responsible for almost all of the PTV association to ASDs^21^ (**Online Methods: De novo *variant analyses***). *De novo* PTVs in this class are a) absent from the Exome Aggregation Consortium database, a reference sample of over 60,000 exomes, and b) found within a gene predicted to be intolerant of heterozygous loss of function variation (probability of loss of function intolerance (pLI) ≥ 0.9)^22^. In the SSC, *de novo* PTVs in this class were found in 7.1% of cases and 2.1% of unaffected siblings (P = 4.12E-14). As *de novo* copy number variants (CNVs) that delete a gene should have the same, if not greater, molecular impact as a PTV in that gene, we were motivated to investigate whether we could similarly refine ASDs’ association with *de novo* deletions (**Online Methods: De novo *variant analyses***). In the SSC, we found that ASDs were strongly associated with *de novo* deletions from two categories: a) deletions that include a constrained gene, as defined above, and b) uncommon, large (≥ 500 kilobase) deletions that do not contain a constrained gene (from here: contributing CNV deletions; **Supplementary Figure 1**). Contributing CNV deletions were seen in 2.4% of SSC cases and 0.5% of SSC unaffected siblings (P = 2.73E-07). All other types of *de novo* deletions were observed in 1.5% of cases and 1.2% of unaffected siblings, and were not associated with ASD risk (P = 0.50; **Supplementary Figure 2**).

Together, these refined classes of *de novo* PTVs and deletions form a category of strong acting *de novo* variants (from here: contributing *de novo* variants (CDNVs); odds ratio (OR) = 3.91, P = 6.56E-20; case rate = 9.4%; control rate = 2.6%). Consistent with a hallmark of ASD-associated *de novo* variation, Figure 3a shows that the rate of CDNVs in SSC varies substantially based on the probands’ counts of co-occurring adverse neurodevelopmental events, in this case the count of: delayed walking, a history of seizures, or intellectual disability (**Supplementary Table 12a**; **Online Methods: De novo *variant analyses***). ASD probands without any of those comorbid traits were more than 3 times (OR = 3.15; P = 3.88E-10) as likely to carry a CDNV than their unaffected siblings; ASD probands with all three of these comorbid traits were approximately 15 times as likely to carry a CDNV than their unaffected siblings (OR = 15.05; P = 9.08E-10). Because *de novo* events observed in cases and controls differ with regard to their average severity, their effect size cannot be directly estimated using the case-control carrier ratio. Using the male:female carrier approach described by de Rubeis et al.^3^, we estimate that, on average, the CDNVs defined here confer an approximate 20-fold increase in risk for an ASD diagnosis (**Online Methods: De novo *variant analyses)***. Their effect size, however, likely varies as the male:female ratio of the carriers declines with increasing neurodevelopmental comorbidities (**Supplementary Table 12b**). In other words, the CDNVs seen in the ASD individuals with multiple comorbid neurodevelopmental traits are likely, on average, more deleterious than those seen in probands with ASD alone.

**Figure 3.**
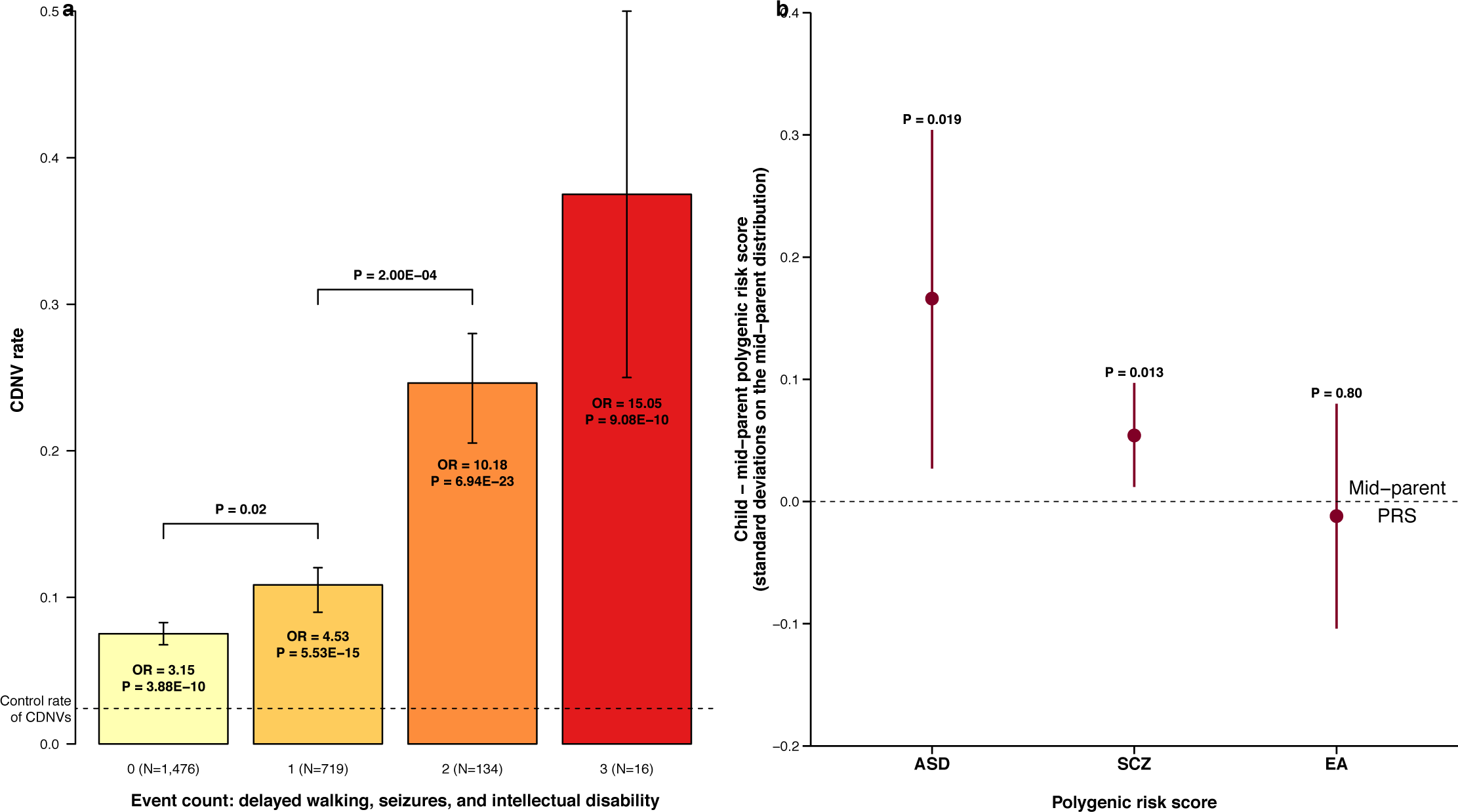
Contributing *de novo* mutations are associated with adverse neurological and developmental events and act additively with polygenic burden to influence ASD risk. (**a**) Simons Simplex Collection (SSC) probands are grouped by their count of the following: delayed walking (≥ 18 months); presence of seizures; intellectual disability (full-scale IQ < 70). Case and control contributing *de novo* variant (CDNV) rate is calculated by dividing the count of CDNVs by the count of individuals in the group. The odds ratio (OR) compares the proband CDNV rate in each event bin to the overall control CDNV rate. P-values in the shaded bars indicate the probability that the CDNV rate in cases is not different from the CDNV rate in controls. (**b**) pTDT analysis for SSC CDNV proband carriers (*n* = 221). Transmission shown in terms of standard deviations on mid-parent distribution ± 1.96 SE (95% confidence intervals). P-values denote the probability that the mean of the pTDT distribution is 0.

Shown in Figure 3b, we used pTDT to determine whether polygenic risk for ASD, SCZ, and EA was also over transmitted to CDNV carriers (**Online Methods: De novo *variant analyses)***. ASD cases with a CDNV (*n* = 221 cases) indeed carried more polygenic risk for ASD and SCZ than expected based on parent background (P < 0.05 for both comparisons). There was no evidence of over transmission of EA PRS (P = 0.80). This provides clear evidence for additivity between common and rare variation in creating risk for ASDs. We did not see a difference in polygenic over transmission between probands with zero versus one or more co-occurring neurodevelopmental events (P > 0.05 for all comparisons; **Supplementary Table 12c**), further supporting the consistent influence of polygenic risk factors.

## Additivity among common polygenic risk factors

In the context of three orthogonal risk distributions, each of which is associated with ASDs, one does not have to maintain an extreme position on any single distribution to carry a cumulatively uncommon amount of ASD risk (Figure 4a). For example, being in the top 20% of three uncorrelated ASD risk distributions would result in a cumulative amount of risk seen in less than 1% of people in the population (0.2^3^=0.008).

**Figure 4.**
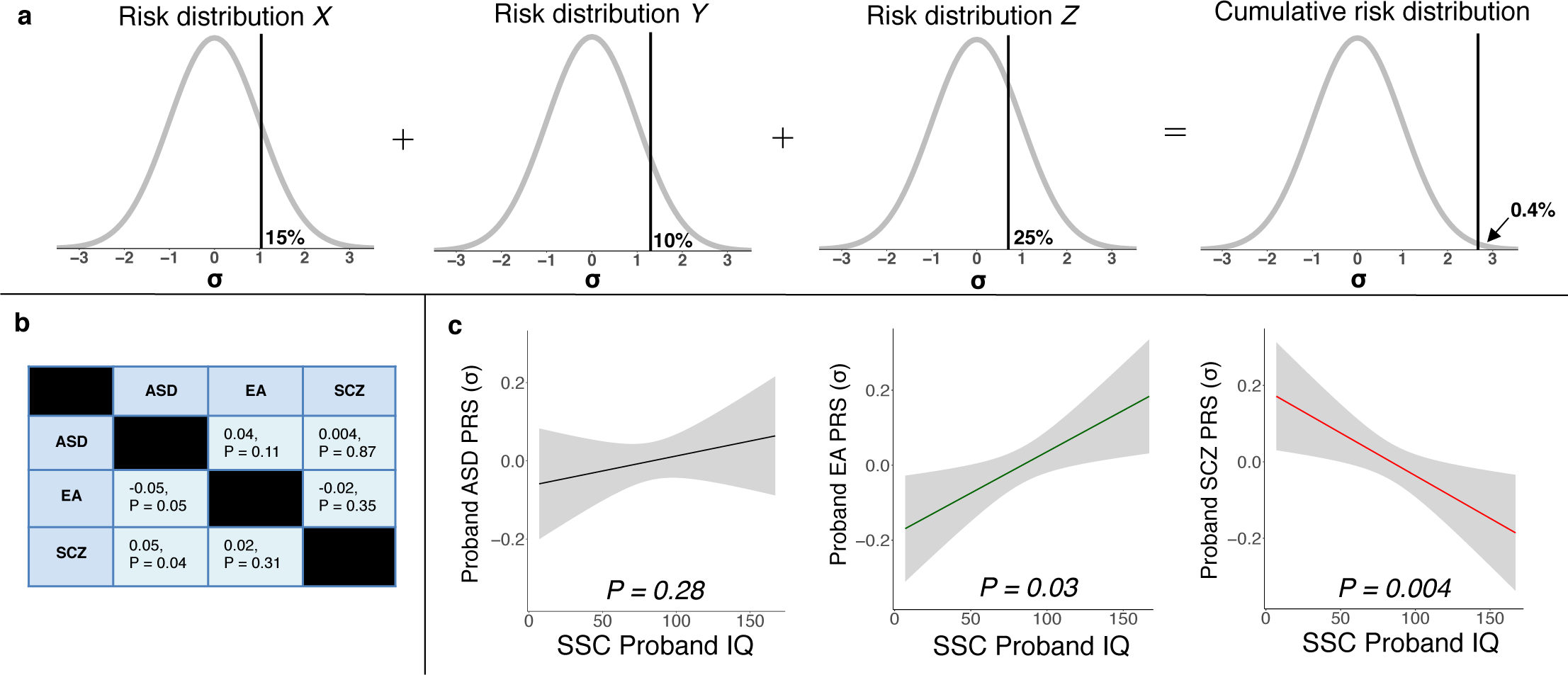
(**a**) Additivity among orthogonal risk factors can yield high cumulative risk. (**b**) Polygenic risk scores (PRS) for ASDs, schizophrenia (SCZ), and educational attainment (EA) are not strongly associated at either the mid-parent level (above diagonal) or the pTDT deviation level (below diagonal). The table contains Pearson correlation coefficients and p-values indicating the probability with which the true correlation is 0. PRS are from European ancestry SSC families. (**c**) Polygenic risk factors for ASD exhibit independent, distinct effects on IQ. P-values indicate the probability with which there is no association between each polygenic risk score (PRS) and SSC proband IQ.

Common polygenic risk for ASDs, SCZ, and EA is partially correlated^6^. As shown in Figure 4b, however, we observed little evidence of correlation between the ASD, SCZ, and EA PRS in terms of either a) correlation at the mid-parent level, above the diagonal or b) correlation in the degree to which the scores were transmitted to the probands (pTDT deviation), below the diagonal. The lack of strong associations is likely a function of both ascertainment effects and attenuation due to limited predictive ability of the PRS.

Given limited association between the three PRS, we were able to examine additivity among largely distinct common polygenic risk factors. First, we saw significant evidence that each of the three PRS were independently over transmitted (P < 2E-04 in all SSC + PGC ASD comparisons**; Online Methods: *Genetic heterogeneity***; **Supplementary Table 13**). This means that ASD risk is influenced by elements unique to each of the scores, as well as by elements that are shared among them. Independent influences are further suggested by the scores’ relationship with proband IQ (**Supplementary Table 14**). In Figure 4c, we show the association between each of the three PRS and full scale proband IQ in European ancestry SSC probands, controlling for sex, presence of CDNVs, the other two PRS, and the first 10 principal components of ancestry (**Online Methods: *Genetic heterogeneity*)**. The EA and SCZ PRS are associated with proband IQ in opposite directions, consistent with the patterns observed in the general population (**Supplementary Table 14**)^14^.

In addition to statistical evidence of unique contributions, the phenotypic associations suggest that the three PRS are influencing ASD risk through at least partially different processes. Polygenic risk for SCZ influences ASD liability in a manner that negatively influences cognition. Polygenic risk for EA influences ASD liability in a manner that positively influences cognition. These findings reinforce the idea that ASD heterogeneity is shaped not only by rare variants of strong effect, but also by common variant risk acting through multiple biological pathways.

Despite longstanding evidence for common polygenic influences on ASD risk, many have questioned those associations, particularly the recently published – and counterintuitive – findings from genetic correlation analysis. Using pTDT, we have shown an unambiguous association between ASD risk and the common polygenic influences on: ASDs themselves, schizophrenia, and greater educational attainment. These effects were evident in affected males and affected females, as well as ASD individuals with and without intellectual disability. Because of the strong association between polygenic predictors of educational attainment and intelligence^14^, this finding means that, on average, individuals with ASD and intellectual disability were genetically predicted to have higher IQ than their parents and unaffected siblings. That association, which replicated in independent ASD cohorts and held in probands without an ASD-associated *de novo* event, will require further study. With regard to proband phenotype, the finding furthers existing questions of whether an IQ measured below 70 in someone with an ASD might be different in some ways from an IQ measured below 70 in someone without. With regard to better understanding the mechanisms through which genetic risk is conferred, we need to examine how and when certain amounts of risk are beneficial (e.g., a strong interest in the arts or sciences that increases one’s educational attainment), and how and when they are deleterious (e.g., an overwhelming interest in a topic that disrupts healthy and necessary activities). Genetic risk and phenotypic traits relevant to neuropsychiatric disease exist on continua^18^, for which effective research and treatment paradigms will likely need to account.

These findings also highlight important differences between the common and rare variant contributors to ASD risk. Strong acting, *de novo* variant risk for ASDs impacts a limited subset of cases. The phenotypic preferences of those types of variants are now well established; they are associated with intellectual disability, seizures, and global neurodevelopmental impact^1^. Common variant risk factors, on the other hand, appear more pervasively influential. Common variant risk appears similarly relevant to ASD individuals with high and low IQ, and with and without a large acting *de novo* mutation. The common polygenic influences also appear comparatively neurologically gentle. They are, in many cases, in fact associated with better educational and cognitive outcomes in the population. These differences strongly suggest that *de novo* and common polygenic variation may confer risk for ASD in different ways. Particularly as common polygenic risk is the more consistent contributor to ASD liability across cases, it will be critically important to take common variation into account in creating animal or stem cell models of ASD.

## Acknowledgements

We thank Drs. Steven Hyman and Rosy Hosking for their thoughtful comments. E.B.R. and D.J.W. were funded by National Institute of Mental Health grant 1K01MH099286-01A1 and Brain Behavior Research Foundation (NARSAD) Young Investigator grant 22379. E.M.W. was funded by the Stanley Center for Psychiatric Research at the Broad Institute. AO was funded by ERC Consolidator Grant (647648 EdGe). We thank the families who took part in the Simons Simplex Collection study and the clinicians who collected data at each of the study sites. The iPSYCH project is funded by the Lundbeck Foundation and the universities and university hospitals of Aarhus and Copenhagen. Genotyping of iPSYCH and PGC samples was supported by grants from the Stanley Foundation, the Simons Foundation (SFARI 311789 to M.J.D.) and the National Institute of Mental Health (5U01MH094432-02 to M.J.D.). This work was also supported by a grant from the Simons Foundation (SFARI 307705; to S.J.S.). The authors would like to thank the Exome Aggregation Consortium and the groups that provided exome variant data for comparison. A full list of contributing groups can be found on the ExAC website (see URLs).

## Online Methods

### Sample Description

The analytic cohorts are presented in **Supplementary Table 1**. The Simons Simplex Collection (SSC) is a resource of more than 2,500 families with a child diagnosed with an autism spectrum disorder (ASD)^19^. Each member of the family was genotyped on one of the following platforms: Illumina Omni2.5, Illumina 1Mv3 or Illumina 1Mv1^1^. We analyzed 2,091 SSC quads, defined as families with both parents, the proband, and a designated unaffected sibling genotyped, and 493 SSC trios, defined as families with both parents and the proband genotyped. Most SSC families were also whole exome sequenced to detect rare coding variation (82.7% of quads, an additional 8.0% of quads without sibling sequencing; 91.1% of trios).

The Psychiatric Genomics Consortium Autism Group (PGC ASD) sample consisted of parent-proband trios from the Psychiatric Genomics Consortium Autism Group (**Supplementary Table 2**). Our PGC ASD analytic sample excluded the SSC families present in the PGC ASD genome-wide association study (GWAS)^23^. The PGC ASD sample described here is accordingly independent from the SSC sample and included 3,870 parent-proband trios. In brief, the PGC ASD data included ASD trio cohorts from: Autism Center of Excellence UCLA (*n* = 215), Autism Genome Project (*n* = 2,254), Montreal/Boston Collection (*n* = 138), Johns Hopkins University (*n* = 764), and the Children’s Hospital of Philadelphia (*n* = 499)^23^. All genotype data (SSC and PGC ASD) were imputed using the 1000 Genomes reference panel and Ricopili pipeline, which are publicly available and have been reported on extensively^20,24,25^.

Though trio approaches are broadly immune to confounding through ancestry, we isolated European-only subsets of both the SSC and PGC ASD samples to a) ensure that our primary results did not change in an ancestrally homogeneous subset of the data and b) conduct comparisons across probands or across parents, which might be sensitive to ancestry (as opposed to comparisons between probands and parents in a trio). In the SSC, we first selected probands with parent-reported white non-Hispanic ancestry (*n* = 1,912). We merged the genotype data from those individuals with the Hapmap III dataset^26^ and generated principal components of ancestry using GCTA^27^. Through visual inspection (**Supplementary Figure 8**), we defined an ancestrally European SSC subcohort, leaving 1,851 probands (and, by extension, 1,851 families). We calculated principal components of ancestry distinctly within the derived SSC European ancestry subcohort and used these as covariates in the non-trio analysis (e.g., genotype to phenotype analyses among probands).

Self-reported ancestry was not available for the PGC ASD samples. To conservatively isolate a European ancestry PGC ASD subcohort, we identified those families in which both parents were of European ancestry. To do so, we merged the PGC ASD parent data with Hapmap III and similarly generated principal components of ancestry using GCTA. By visual inspection, we identified European ancestry individuals, leaving 6,742 of the original 7,740 PGC ASD parents (**Supplementary Figure 9**). Both parents were of European ancestry in 3,209 families, and those families comprised the European ancestry PGC ASD subcohort. We again calculated principal components of ancestry within the derived PGC ASD European ancestry subcohort for use as analytic covariates in non-trio analyses.

### Polygenic Risk Scoring

A polygenic risk score (PRS) provides a quantitative estimate of an individual’s genetic predisposition (“risk”) for a given phenotype based on common variant genotype data and independent GWAS results for the target phenotype (e.g., schizophrenia or educational attainment)^28^. The score provides a relative, not absolute, measure of risk. For example, individual A (PRS_schizophrenia_ = 8) is at higher estimated genetic risk for schizophrenia than individual B (PRS_schizophrenia_ = 6), but a PRS of 8 or 6 is not independently interpretable.

We calculated polygenic risk for ASD, educational attainment (EA), schizophrenia (SCZ) and body mass index (BMI) for all individuals with available genotype data in the SSC and PGC ASD datasets. To do so, we used summary statistics from the largest available, independent GWAS of each phenotype (**Supplementary Table 3**). None of the subjects in the SSC or PGC ASD cohort were included in any of the four GWAS discovery samples. We selected SCZ and EA because of their robust, well-replicated associations with ASD risk^6,14,23^. We selected BMI as a negative control due to its lack of association with ASD risk^6^. We used ASD summary statistics from a GWAS of a Danish population-based sample of 7,783 cases and 11,359 controls from the first 10 genotyping waves of the iPSYCH-Broad Autism project^18^. The SCZ summary statistics were from the 2014 GWAS of 36,989 cases and 113,075 controls from the Psychiatric Genomics Consortium^20^. The EA summary statistics were from a population-based GWAS of years of schooling (*n* = 328,917, discovery and replication meta-analysis, excluding 23andMe)^12^. The BMI summary statistics were from a population-based GWAS of body mass index (*n* = 322,154, European ancestry meta-analysis)^29^.

To construct the polygenic risk scores, we first gathered the summary statistics from the GWAS described above. The summary statistics included effect sizes and p-values for each single nucleotide polymorphism (SNP) in the GWAS. We employed the widely used Ricopili pipeline to generate the PRS. In brief, we removed SNPs within 500 kilobases of and correlated (r^2^ ≥ 0.1) with a more significantly associated SNP in the GWAS^20^. We used the 1000 Genomes Reference panel to estimate SNP correlations^24^. For complex, polygenic traits, only a small fraction of SNPs pass the genome-wide significance threshold of P = 5.00E-08; the majority of the signal resides in SNPs that are less significantly associated^6,12,20,27^. In order to maximally capture common, polygenic influence, we relaxed the p-value threshold for SNPs in the PRS until doing so added more noise than signal (threshold options: P ≤ 1, 5E-1, 2E-1, 1E-1, 5E-2, 1E-2, 1E-3, 1E-4, 1E-6, and 5E-8). We identified the optimal p-value threshold as that which explained the most phenotypic variation for each trait in an independent sample. For the ASD PRS, we found that the threshold of P = 0.1 explained the most case-pseudocontrol variance in SSC (as the SSC does not have independent control data, we generated pseudocontrol genotypes from the untransmitted parental alleles)^18,23^. For the SCZ and EA PRS, we used the threshold identified by the discovery GWAS analyses as explaining the most variance. In the PGC’s 2014 schizophrenia analyses, a p-value threshold of P = 0.05 most commonly explained the most SCZ case-control variance in 40 leave-one-out analyses^20^. For educational attainment, constructing a PRS using a threshold of P = 1 explained the most variance in number years of education achieved in an independent sample^12^. For BMI, constructing a PRS using a threshold of P = 0.2 explained the most variance in phenotypic body mass index in the Simons Simplex Collection (**Supplementary Methods 9**). These thresholds create the strongest polygenic risk scores for ASD, EA, SCZ and BMI, leaving us maximally powered to investigate the relationship between these four traits and ASDs.

Next, we excluded SNPs that were poorly imputed in either SSC or PGC ASD (info score < 0.6 in either cohort). The exception to this filtering rule was in SSC-specific analyses, (e.g., analysis of *de novo* variation) where our info threshold of 0.6 was the minimum in the SSC imputation only. For each trait, the polygenic risk scores for individuals in SSC and PGC ASD were then calculated as the product of the GWAS effect size (log odds or beta) at that SNP by the individual’s count of reference alleles at that SNP (0, 1 or 2), summed across all remaining markers. We implemented this scoring protocol using the score function in Plink^30^. If an individual was missing genetic data at a SNP in the summary statistics file, Plink calculated the expected score based on the cohort-wide allele frequency.

### pTDT

We define pTDT deviation as:

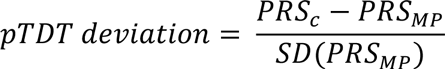

where PRS_C_ is the polygenic risk score for the child (proband or unaffected sibling). PRS_MP_ is the mid-parent polygenic risk score:

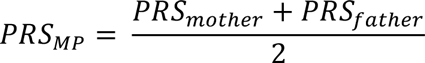

SD(PRS_MP_) is the standard deviation of the sample-specific mid-parent PRS. For example, in SSC analysis, the SD(PRS_MP_) was the standard deviation of the mid-parent PRS distribution in SSC parents. The approach can be adapted to PRS from unaffected siblings, but using mid-parent PRS improves statistical power (**Supplementary Method 4, Supplementary Table 17**).

To evaluate whether the pTDT deviation is significantly different than 0, we defined the pTDT test statistic (*t_pTDT_*) as:

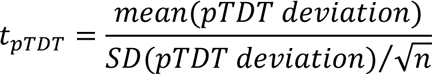

where *n* is the number of families included in the pTDT. We evaluate t_pTDT_ as a one-sample t-test.

We performed pTDT using the ASD, EA, SCZ and BMI PRS described above in four groups: SSC probands (*n* = 2,584), SSC unaffected siblings (*n* = 2,091), PGC ASD probands (*n* = 3,870), and the combination of SSC and PGC ASD probands (*n* = 6,454). As described in the main text (Figure 2a), each of the ASD, EA, and SCZ PRS were significantly over transmitted from parents to probands, but not to unaffected siblings (P < 1E-06 for all parent to proband comparisons in either SSC or PGC ASD; P < 1E-15 for all parent to proband comparisons in combined SSC and PGC ASD; P > 0.05 for all unaffected sibling comparisons). In contrast, BMI PRS was not over transmitted to probands (P > 0.05). In a complementary permutation test, we randomly assigned case/control labels to the affected and unaffected children in SSC. We then counted the number of times that the simulated difference in ASD PRS between affected and unaffected SSC children exceeded the observed difference (86 of 1,000,000 permutations = 0.0086%), consistent with the primary results (Figure 2a).

In our primary associations (Figure 2a, **Supplementary Table 7a**), there were no statistically significant pTDT differences between the SSC and PGC ASD cohorts (**Supplementary Table 7b**). We accordingly combined SSC and PGC ASD for the analyses stratified by sex and IQ to increase statistical power (Figures 2b, 2c). In SSC, full scale IQ (FSIQ; SSC variable: sscfsiq) was derived from a number of scales and available for almost all probands (99.8%) (Differential Ability Scales, Second Edition^31^; Mullen Scales of Early Learning^32^; Wechsler battery^33^). Our full scale SSC IQ estimates were taken from the SSC’s ‘full scale deviation IQ’ variable when it was available, and ‘full scale ratio IQ’ when it was not^34^. FSIQ was measured heterogeneously across the contributing PGC ASD cohorts (**Supplementary Table 2**). To accommodate measurement differences, FSIQ was converted to broad groups for PGC consortium-level analyses (for the most part, numeric IQ values from PGC ASD were not available for this analysis). PGC ASD probands were assigned FSIQ levels 1-4 as follows: (1) FSIQ < 35; (2) 35 ≤ FSIQ < 70; (3) 70 ≤ FSIQ < 90; (4) 90 ≤ FSIQ. In PGC ASD, 38.9% of probands had estimated IQ available. We divided each of SSC and PGC ASD by presence/absence of intellectual disability (ID) in the proband so that they could be analyzed together (SSC ID: IQ < 70; PGC ASD ID: IQ = 1 or IQ = 2). We repeated the IQ stratified analyses in SSC and PGC ASD separately, which further suggested no differences between the cohorts, despite the limited FSIQ data available in PGC ASD (**Supplementary Tables 9a and 9b**).

### *De novo* variant analyses

We defined a group of *de novo* mutations strongly associated with ASD risk (**Supplementary Table 11**). We performed this analysis exclusively in SSC; we could not perform the analysis in PGC ASD because only common variant (GWAS) data was available. As detailed in the main text, our previous work has identified a subclass of *de novo* protein truncating variants (PTVs) that are a primary source of association to ASD^21^. PTVs not in this contributing class are not associated with proband IQ (P = 0.67) (**Supplementary Figure 6**) and are unassociated with ASD risk (observed in 7.8% of cases and 6.9% of unaffected siblings, P = 0.33).

As detailed in the main text, *de novo* copy number variants (CNVs) that delete a gene should have the same molecular impact as a PTV in that same gene. The associations between CNV categories and ASD risk are presented in the supplement and did not differ substantially when controlling for parental age at birth of proband (**Supplementary Table 16**). Contributing deletions outside of our defined class were not associated with proband IQ (P = 0.34, **Supplementary Figure 4**). In contrast, *de novo* duplications of constrained genes are not disproportionately associated with ASD risk (P = 0.49; **Supplementary Figure 3**, **Supplementary Methods 5**).

We identified a subset of *de novo* CNV deletions, primarily those containing loss of function intolerant genes, that account for most of the category’s association to ASDs (contributing CNV deletions; **Supplementary Figure 1**). Together, contributing PTVs and contributing CNV deletions formed a strong acting *de novo* variant category in SSC, known from here as contributing *de novo* variants (CDNVs). Multiple lines of evidence suggest CDNVs are robustly associated with ASD risk. CDNVs were seen in 9.4% of SSC cases (*n* = 221 of 2,346) that were both sequenced and genotyped and 2.6% of SSC unaffected siblings (*n* = 45 of 1,736) that were both sequenced and genotyped (P = 6.56E-20). CDNVs were very strongly associated with proband IQ (P = 5.92E-06; controlling for proband sex), while all other *de novo* PTVs and deletions (those not in the CDNV category) were not associated with proband IQ (P = 0.44, controlling for proband sex; **Supplementary Figure 5**).

As *de novo* variants are on average more severe when observed in ASD cases than in controls^35^, we could not estimate the amount of ASD risk conferred by CDNVs directly from the case-unaffected sibling odds ratio. As noted in the main text, we used the male:female CDNV carrier ratio to re-estimate the effect size for the event class, as described in De Rubeis et al., 2014. In brief, the approximately 4:1 male:female ratio among ASD cases suggests a different ASD liability threshold for males and females in the population. Regardless of whether that difference reflects etiology or ascertainment, it results in a direct mathematical relationship between the expected effect size of an event class and the male:female carrier ratio of cases. In the case of CDNVs, a variant class observed twice as frequently in female probands than in male probands (17.4% v 8.2%; P = 1.35E-06; **Supplementary Figure 7**), we estimate an OR of approximately 20^3^. This estimate far exceeds that directly suggested by the CDNV case-control excess of 3.63.

We next examined the rate of CDNVs in SSC probands based on the number of adverse proband co-occurring neurological and developmental events (Figure 3a). Previous studies have demonstrated that *de novo* PTVs and *de novo* CNVs are both associated with intellectual disability (IQ < 70) in ASD probands and positively associated with history of seizures^1,15^. Motor delay is also a neurodevelopmental outcome associated with autism spectrum disorder^36^. Here, we defined motor delay for SSC probands as failing to walk unaided before 18 months after birth, the age by which the great majority of children have begun to walk^37^. As hypothesized, motor delay, seizures, and low IQ were independently associated with CDNV rate in SSC probands after controlling for proband sex (**Supplementary Table 12a**). CDNV rate was positively associated with the number of these adverse events experienced by probands (Figure 3a, **Supplementary Table 12b**). The decreasing male:female ratio among CDNV carriers as count of co-occurring neurodevelopmental outcomes increases suggests that the co-occurring neurodevelopmental categories are capturing a severe phenotype (**Supplementary Method 6**, **Supplementary Table 12b**). Multiple lines of evidence, including associations with IQ, male:female carrier ratio, and adverse co-occurring neurodevelopmental events, suggest that CDNVs are strongly associated with ASD risk.

Using pTDT, we evaluated polygenic transmission in probands carrying at least one CDNV (Figure 3b, **Supplementary Table 10**). As this CDNV analysis is specific to SSC, the polygenic risk scores generated for the CDNV analysis used info score cutoffs from SSC imputation in order to increase the number of well-imputed SNPs retained for PRS. We restricted our analysis to those families with genotyped parents and probands with both genotype and exome sequence data (*n* = 2,345; *n* = 221 with CDNV). The cohort of probands with a CDNV is too small to determine whether the difference in transmission between probands that do and do not carry a CDNV is statistically significant. We also analyzed whether polygenic over transmission was seen in a more broadly defined set of *de novo* events (**Supplementary Methods 7, Supplementary Table 10**).

### Genetic heterogeneity and proband phenotype

We analyzed whether polygenic risk for ASD, EA and SCZ were each independently over transmitted to ASD cases. For each of SSC (*n* = 2,584 trios), PGC ASD (*n* = 3,870 trios) and SSC & PGC ASD combined (*n* = 6,454 trios), we performed a logistic regression predicting proband/average parent status from polygenic risk for ASD, EA and SCZ. We confirmed that the over transmission acts independently in each PRS (P < 2E-04 for all PRS in SSC + PGC ASD cohort; **Supplementary Table 13**).

To calculate the relationship between proband polygenic risk, CDNVs and ASD IQ, we performed a linear regression using proband sex, the three polygenic risk scores and CDNV count to predict full-scale IQ of SSC probands of European ancestry. The results were consistent with previously published association between the three polygenic risk scores and IQ in the general population (**Supplementary Table 14)**^14^. This relationship between polygenic risk and proband IQ holds when using mid-parent PRS and controlling for proband CDNV status (**Supplementary Methods 8, Supplementary Table 15**).

## References

1. Sanders, S.J. et al. Insights into Autism Spectrum Disorder Genomic Architecture and Biology from 71 Risk Loci. Neuron 87, 1215–33 (2015).

2. Gaugler, T. et al. Most genetic risk for autism resides with common variation. Nat Genet 46, 881–5 (2014).

3. De Rubeis, S. et al. Synaptic, transcriptional and chromatin genes disrupted in autism. Nature 515, 209–15 (2014).

4. Iossifov, I. et al. The contribution of de novo coding mutations to autism spectrum disorder. Nature 515, 216–21 (2014).

5. Bulik-Sullivan, B.K. et al. LD Score regression distinguishes confounding from polygenicity in genome-wide association studies. Nat Genet 47, 291–5 (2015).

6. Bulik-Sullivan, B. et al. An atlas of genetic correlations across human diseases and traits. Nat Genet 47, 1236–41 (2015).

7. Krumm, N. et al. Excess of rare, inherited truncating mutations in autism. Nat Genet 47, 582–8 (2015).

8. Anney, R. et al. Individual common variants exert weak effects on the risk for autism spectrum disorders. Hum Mol Genet 21, 4781–92 (2012).

9. Klei, L. et al. Common genetic variants, acting additively, are a major source of risk for autism. Mol Autism 3, 9 (2012).

10. Onis, M. WHO Motor Development Study: Windows of achievement for six gross motor development milestones. Acta Paediatrica 95, 86–95 (2006).

11. Deciphering Developmental Disorders, S. Large-scale discovery of novel genetic causes of developmental disorders. Nature 519, 223–8 (2015).

12. Okbay, A. et al. Genome-wide association study identifies 74 loci associated with educational attainment. Nature 533, 539–42 (2016).

13. Clarke, T.K. et al. Common polygenic risk for autism spectrum disorder (ASD) is associated with cognitive ability in the general population. Mol Psychiatry (2015).

14. Hagenaars, S.P. et al. Shared genetic aetiology between cognitive functions and physical and mental health in UK Biobank (N=112 151) and 24 GWAS consortia. Mol Psychiatry (2016).

15. Robinson, E.B. et al. Autism spectrum disorder severity reflects the average contribution of de novo and familial influences. Proc Natl Acad Sci U S A 111, 15161–5 (2014).

16. Spielman, R.S., McGinnis, R.E. & Ewens, W.J. Transmission test for linkage disequilibrium: the insulin gene region and insulin-dependent diabetes mellitus (IDDM). Am J Hum Genet 52, 506–16 (1993).

17. Wright, C.M. & Cheetham, T.D. The strengths and limitations of parental heights as a predictor of attained height. Arch Dis Child 81, 257–60 (1999).

18. Robinson, E.B. et al. Genetic risk for autism spectrum disorders and neuropsychiatric variation in the general population. Nat Genet 48, 552–5 (2016).

19. Fischbach, G.D. & Lord, C. The Simons Simplex Collection: a resource for identification of autism genetic risk factors. Neuron 68, 192–5 (2010).

20. Schizophrenia Working Group of the Psychiatric Genomics, C. Biological insights from 108 schizophrenia-associated genetic loci. Nature 511, 421–7 (2014).

21. Kosmicki, J. et al. Refining the role of de novo protein truncating variants in neurodevelopmental disorders using population reference samples. bioRxiv (2016).

22. Lek, M. et al. Analysis of protein-coding genetic variation in 60,706 humans. Nature 536, 285–91 (2016).

23. Cross-Disorder Group of the Psychiatric Genomics, C. et al. Genetic relationship between five psychiatric disorders estimated from genome-wide SNPs. Nat Genet 45, 984–94 (2013).

24. Genomes Project, C. et al. An integrated map of genetic variation from 1,092 human genomes. Nature 491, 56–65 (2012).

25. Ricopili. Rapid Imputation Consortium Pipeline.

26. International HapMap, C. et al. Integrating common and rare genetic variation in diverse human populations. Nature 467, 52–8 (2010).

27. Yang, J., Lee, S.H., Goddard, M.E. & Visscher, P.M. GCTA: a tool for genomewide complex trait analysis. Am J Hum Genet 88, 76–82 (2011).

28. Wray, N.R., Goddard, M.E. & Visscher, P.M. Prediction of individual genetic risk to disease from genome-wide association studies. Genome Res 17, 1520–8 (2007).

29. Locke, A.E. et al. Genetic studies of body mass index yield new insights for obesity biology. Nature 518, 197–206 (2015).

30. Chang, C.C. et al. Second-generation PLINK: rising to the challenge of larger and richer datasets. Gigascience 4, 7 (2015).

31. Elliott, C. Differential Ability Scales, (The Psychological Corporation, San Antonio, TX, 2007).

32. Mullen, E. Mullen Scales of Early Learning, (American Guidance Service, Circle Pines, MN, 1995).

33. Wechsler, D. Wechsler abbreviated scale of intelligence, (Psychological Corporation, San Antonio, TX, 1999).

34. Chaste, P. et al. A genome-wide association study of autism using the Simons Simplex Collection: Does reducing phenotypic heterogeneity in autism increase genetic homogeneity? Biol Psychiatry 77, 775–84 (2015).

35. Samocha, K.E. et al. A framework for the interpretation of de novo mutation in human disease. Nat Genet 46, 944–50 (2014).

36. Provost, B., Lopez, B.R. & Heimerl, S. A comparison of motor delays in young children: autism spectrum disorder, developmental delay, and developmental concerns. J Autism Dev Disord 37, 321–8 (2007).

37. World Health Organization, T. WHO Motor Development Study: windows of achievement for six gross motor development milestones. Acta Paediatr Suppl 450, 86–95 (2006).

